# Vaccine- and BTI-elicited pre-Omicron immunity more effectively neutralizes Omicron sublineages BA.1, BA.2, BA.4 and BA.5 than pre-Omicron infection alone

**DOI:** 10.1101/2022.10.24.513415

**Authors:** Eveline Santos da Silva, Jean-Yves Servais, Michel Kohnen, Victor Arendt, Georges Gilson, Therese Staub, Carole Seguin-Devaux, Danielle Perez-Bercoff

## Abstract

Since the emergence of SARS-CoV-2 Omicron BA.1 and BA.2, several Omicron sublineages have emerged, supplanting their predecessors. BA.5 is the current dominant sublineage. Here we compared the neutralization of Omicron sublineages BA.1, BA.2, BA.4 and BA.5 by human sera collected from individuals who were infected with the ancestral B.1 (D614G) strain, vaccinated (3 doses), or with hybrid immunity from vaccination (2 doses) followed by pre-Omicron breakthrough infection (BTI) with Gamma or Delta. All Omicron sublineages exhibited extensive escape from all sera compared to the ancestral B.1 strain and to Delta, albeit to different levels depending on the origin of the sera. Convalescent sera were unable to neutralize BA.1, and partly neutralized BA.2, BA.4 and BA.5. Vaccinee sera partly neutralized BA.2, but BA.1, BA.4 and BA.5 evaded neutralizing antibodies. BTI sera were either non-neutralizing or partially neutralizing. In this case, they had similar neutralizing ability against all Omicron sublineages. Despite similar levels of anti-Spike and anti-Receptor Binding Domain (RBD) antibody in all groups, BTI sera had the highest cross-neutralizing ability against all Omicron sublineages and convalescent sera were the least neutralizing. The NT50:antibody titer ratio, which reflects antibody avidity, was significantly higher in sera from BTI patients compared to convalescent sera, underscoring qualitative differences in antibodies elicited by infection alone and by vaccination. Together these findings highlight the importance of vaccination to trigger highly cross-reactive antibodies that neutralize phylogenetically and antigenically distant strains, and suggest that immune imprinting by first generation vaccines may restrict, but not abolish cross-neutralization.

## Introduction

The Omicron lineage of SARS-CoV-2 comprises several sublineages. BA.1, BA.2 and BA.3 were first identifed in South Africa in November 2021. Between December 2021 and early January 2022, Omicron BA.1 rapidly outcompeted the Delta variant, which dominated the COVID-19 landscape across all continents at the time (1). BA.1 was rapidly followed by the genetically distinct BA.2 sublineage, generating two overlapping peaks in most countries, including Luxembourg, in the winter and early spring 2022. BA.1 and BA.2 harbor 29 and 34 mutations, insertions and deletions in Spike respectively, of which 21 are shared by the two sublineages. BA.3 shares mutations with both BA.1 and BA.2 but has spread poorly compared to BA.1 and BA.2. Within a few months, other BA.2-derived sublineages, such as BA.4, BA.5 and BA.2.12.1 displaced BA.1 and BA.2. BA.4 and BA.5 have identical Spike sequences and differ by only three mutations in ORF7b, M (Membrane protein) and N (Nucleocapsid). Nevertheless, BA.5 has become the dominant variant in most countries. Since the summer 2022, the Omicron landscape is expanding further. BA.2 sublineages BA.2.12.1, BA.2.75, BA.2.75.2, as well as BA.4, BA.5 and their offspring BA.4.6 and BQ.1.1 are gaining ground. These strains carry additional mutations which further increase their infectivity and antibody escape compared to the parental BA.2 (2–7).

While Omicron strains are less pathogenic and associated with lower fatality rates than Delta, the later strains BA.2.75 and BA.5 seem to gain pathogenicity compared to the early BA.1 and BA.2 (6, 8–18). The Omicron lineage typically features strongly enhanced transmissibility compared to Delta and pre-Omicron variants (3–6, 19, 20). Increased transmissibility is due to its stronger docking to the receptor ACE2, and to its endocytosis-mediated, TMPRSS-2 independent entry into target cells, which also favors immune evasion (5, 11, 16, 21–29), although BA.5 can use the TMPRSS2 route (20). Furthermore, compensatory mutations have appeared in a number of strains to balance the loss replicative cost imposed by immune escape mutations (e.g.R493Q for F486V in BA.4/5 (2, 30, 31)). The Omicron Spike also adopts a distinct, more compact conformation and glycosylation patterns which shield it from type 1, 2 and 3 Neutralizing antibodies (NAbs) (29, 32, 33).

Typically, BA.1 and BA.2 show a dramatic drop in susceptibility to neutralization compared to the ancestral B.1 strain containing the Spike D614G mutation (21–24, 34–46). Sera from vaccinees who have received 2 vaccine doses do not cross-neutralize Omicron sublineages BA.1 and BA.2 (4, 21–24, 30, 33, 38, 40, 44, 45, 47, 48). Vaccination-induced as well as infection-induced NAbs wane after a few months, even after the 3^rd^ or booster dose. Booster vaccination (3^rd^ and 4^th^ doses) effectively restore NAbs and cellular responses against Omicron variants and protect against severe COVID-19 and death (24, 25, 30, 40, 43, 44, 48–61), but their durability is short (16, 33, 60, 61). BA.2 offspring BA.2.12.1 and BA.2.75 are modestly more resistant to NAbs than BA.2, while BA.2.75.2, BA.4 and BA.5 are typically more resistant to vaccine-elicited antibodies than BA.2 due to mutations such as L542R and F486V/S (2, 4, 16, 30, 31, 50, 61–64).

BA.2.75, BA.4 and BA.5 also resist antibodies elicited by Omicron BA.1 and BA.2 infection (2, 3, 16, 30, 31, 50, 63–66).

The emergence of antigenically distinct variants with increased infectivity and ability to evade immune responses elicited by prior infection or vaccination shapes both the pandemic landscape and clinical burden (2, 63, 67). With the exponential increase in breakthrough infections due to Omicron, numerous studies have investigated cross-protection between Omicron sublineages. Epidemiological studies suggest that breakthrough infection with BA.1 and BA.2 confers some degree of cross-protection against infection with other Omicron sublineages such as BA.2-offspring (68, 69). This better protection may reflect better cross-neutralization due to antigenically closer strains, or the shorter time elapsed since infection (19, 64, 69–71). Aside from vaccine-induced immunity, the cross-neutralization by pre-Omicron elicited antibodies has been less studied (38, 41, 42, 57, 72). Yet, it is acknowledged that the first encounter with an antigen molds the immune response and this ‘immune imprinting’, also known as ‘original antigenic sin’ may limit and compromise the subsequent immune response. Depending on the original and challenge antigens, immune imprinting can be helpful if reactivation of existing memory B-cells rapidly provides antibodies that at least partially neutralize the virus, while new, variant-adapted NAbs are generated. Immune imprinting can however obstruct the generation of new, better adapted antibodies, either by neutralizing the antigen which is supposed to boost immunity (thus decreasing the impact of booster doses), or by skewing immunity to continue to produce antibodies against the past virus, impairing the generation of antibodies better suited to neutralize the new variant (2, 63, 72, 73). Imprinting has been beneficial up to Delta, but the completely different Omicron lineages evade antibodies and T-cells elicited by prior immunogens (2, 3, 20, 34, 66, 72–74). Conversely, studies on the cross-protection elicited by Omicron BA.1 infection alone against pre-Omicron and other Omicron sublineages consistently document poor cross-neutralization, highlighting the strong immune imprinting of this sublineage (3, 20, 30, 42, 50, 63, 66, 73, 75, 76). Obstructive immune imprinting has been described for BA.1, BA.2 and BA.5 breakthrough infections after vaccination with first generation vaccines as well (2, 18, 63, 73, 74). In contrast, BA.2-elicited immunity confers cross-protection against BA.5 (16, 64, 77) and BA.2-offspring (16, 64, 77), emphasizing the beneficial impact of reactivation of NAbs targeting common epitopes. Importantly however, mixed immunity resulting from Omicron BA.1-and BA-2-breakthrough infections after 2 or 3 first generation vaccine doses or from reinfection after a first infection with a pre-Omicron strain or from bivalent vaccines combining wild-type + Omicron BA.1, wildtype + Omicron BA.4/5, Delta + Omicron BA.1 or hybrid Omicron BA.1/Delta determinants, all elicit superior NAb as well as T-cell responses against all Omicron sublineages and against pre-Omicron VOCs and confer better protection against severe forms of COVID-19 compared to boosters based on the original Wuhan strain or pre-Omicron (Beta or Delta) (3, 5, 16, 20, 38, 42, 53, 66, 67, 73, 75, 76, 78, 79). Accordingly, second generation bivalent vaccines combining ancestral and Omicron sequences (generally BA.1 or BA.4/BA.5) have been approved by the EMA and FDA in September 2022 as boosters after first generation vaccines based on the original Wuhan strain.

In this context, and as BA.2 and its sublineages BA.4 and BA.5 continue to evolve, it is essential to have a clear view on the cross-neutralization of immune responses induced by infection, by vaccination and by both (hybrid immunity). In this study, we focused on pre-Omicron immunity. We compared the ability of sera from 58 individuals infected with the ancestral B.1 strain before vaccination (convalescent sera), 14 triple-vaccinated individuals and 16 breakthrough infection patients (BTI) infected with Delta (14 patients) or Gamma (2 patients) to neutralize the pre-Omicron strains B.1 (D614G) and Delta and the four main Omicron sublineages BA.1, BA.2, BA.4 and BA.5. We show that convalescent sera had the lowest neutralizing ability and BTI sera had the highest neutralizing ability against all strains. Convalescent sera from patients with moderate disease had better neutralizing ability. Overall, BA.1 was the most resistant to neutralization by all sera, while BA.2 remained the most sensitive to all sera and BA.4 and BA.5 had intermediate resistance levels. However, sera from convalescent, vaccinee and BTI patients showed specificities in their ability to neutralize BA.5, which escaped neutralization by vaccinees better than infection-induced and hybrid immunity. Antibody avidity estimated from the NT50:antibody titer was also highest in BTI, providing some insight into the better efficacy of hybrid immunity.

## Materials and Methods

### Patient/donor samples

This study included sera from 58 unvaccinated patients (hereafter ‘convalescent sera’) infected between March and July 2020, sera from 14 individuals who received a mRNA booster dose between October 2021 and January 2022) and sera from 16 vaccinated patients with breakthrough infection (2 Gamma and 14 Delta) (hereafter ‘BTI sera’) who were infected between July 15^th^ and September 20^th^ 2021. All infected patients had RT-PCR-confirmed SARS-CoV-2 infection. The study was approved by the LIH Institutional Review Board (study number 14718697-NeutraCoV) and was performed in accordance with the 2018 Helsinki Declaration. Anonymized patient left-over samples collected at the Centre Hospitalier de Luxembourg (CHL) were used for the set-up of serological and virological tests in agreement with GDPR guidelines. No clinical data was available for any of the donors. The only available data for the unvaccinated patients was disease severity recorded by the clinician. Disease severity stratification was as follows: *Mild/asymptomatic* patients (7 patients) presented flu-like symptoms or no symptoms; patients with *Moderate* disease (17 patients) had fever, flu-like symptoms, anosmia, fatigue, gastro-intestinal disturbances, but did not require hospitalization or oxygen supplementation; patients with *severe* or *critical* disease (34 patients) were admitted to the hospital, required oxygen supplementation and/or intensive care. Convalescent sera were collected during acute infection (median 16.7 days after symptom onset, interquartile range [IQR] = 13.53-19.87 days), while the BTI sera were collected at the time of diagnosis but time since symptom onset was not known. For BTI cases, data on the lineage of the infecting strain and the time since 2^nd^ vaccine dose were provided by CHL. For vaccinees, only the date of booster dose was available. Median time elapsed since booster (3^rd^) dose was 4 months (IQR = 2.26 – 6 months).

### Cells

Vero-E6 cells (a kind gift from Dr. Thorsten Wolff, Influenza und respiratorische Viren, Robert Koch Institute, Germany) were maintained in DMEM supplemented with 10% Foetal Bovine Serum (FBS), 2 mM L-Glutamine, 50 μg/mL Penicillin and 50 μg/mL Streptomycin (all from Invitrogen, Merelbeke, Belgium). For infection experiments, 2% FBS was used (hereafter referred to as Viral Growth Medium, VGM). HEK293T were from ATCC and were maintained in DMEM supplemented with 10% FBS, 50 μg/mL Penicillin and 50 μg/mL Streptomycin. HEK293T-ACE2-TMPRSS2 cells (SL222 HEK293T,GeneCopeia via Labomics, Nivelles, Belgium). They were maintained in HEK medium containing Puromycin (1 μg/mL) and Hygromycin (100 μg/mL).

### Serology

The MesoScale Diagnostics (MSD) V-Plex COVID-19 Coronavirus Panel 1 serology kit (K15362U) was used according to the manufacturer’s recommendations to determine the IgG profile of sera (MesoScale Diagnostics, Rockville, MD, USA). This multiplex assay includes SARS-CoV-2 antigens (N, S, RBD, NTD) as well as Spike proteins from other Coronaviruses (SARS-CoV, MERS-CoV, OC43, HKU1) and Influenza A Hemagglutinin H3.

### Virus isolation and titration

SARS-CoV-2 strains D614G and VOCs (Gamma, Delta and Omicron) were isolated from anonymized left-over patient nasopharyngeal swabs (NPS) collected from patients at the CHL and the Laboratoire Nationale de Santé to set-up the virological assays. For isolation, 500 μl of residual swab preservation medium was added to Vero-E6 cells (1.2×10^6^ cells) in VGM and cytopathic effect (CPE) was monitored visually daily. Viral supernatant was used to constitute a viral stock by infecting Vero-E6 cells in a second passage. The viral supernatant from passage 2 was centrifuged and stored at −80°C until use. All experiments were performed with the same viral stock. Viral strains present in the original material (swabs) were identified through next-generation sequencing and the Spike was resequenced after the second passage to verify sequence conservation. We isolated representative strains for B.1 (D614G, pre-VOC), Gamma, Delta and Omicron (sublineages BA.1, BA.2, BA.4 and BA.5).

The 50% Tissue Culture Infectious Dose (TCID50) was assessed by titrating viral strains on Vero-E6 cells in sextuplicate wells as previously described (46). Briefly, 10^4^ cells/well were infected with 200 μl of serial 10-fold dilutions of isolated virus (starting at 1:100 in VGM) for 72 hours at 37°C with 5% CO_2_. Virus-induced CPE was measured using the tetrazolium salt WST-8, which is cleaved to a soluble strongly pigmented formazan product by metabolically active cells (CCK-8 kit, Tebu-Bio, Antwerp, Belgium). Optical density at 570 nm was then measured. Virus-exposed wells were compared to uninfected wells (100% survival). The threshold for infection was set at 75% cell survival (i.e. all virus-exposed wells with < 75% viable cells were considered infected) based on preliminary comparative experiments with visually recorded CPE and crystal violet staining. The TCID50 was calculated according to the Reed and Muench method.

### Live-virus neutralization Assay

The live-virus neutralization assay has been described previously (46). Briefly, serial two-fold dilutions of heat-inactivated (30 min at 56°C) patient serum were incubated 1 hour with 100 TCID50 of virus in VGM. The mixture (200 μl/well) was then inoculated on Vero-E6 cells (10^4^ cells/well in a 96-well microtitre plate) and cells were cultured for another 72 hours at 37°C with 5% CO_2_. A positive control (no serum) and an uninfected control (no serum - no virus) were included in each plate to assess maximum infection (no serum) and minimum (no virus) values. All infections were performed in triplicate wells. Virus-induced CPE was measured using the tetrazolium salt WST-8 as above. Percent survival was calculated relative to uninfected cells. The half-maximal inhibitory concentration for serum (IC50) was determined by inferring the 4-parameter non linear regression curve fit (GraphPad Prism v5) with unconstrained top and bottom values. The IC50 was log-transformed into 50% neutralizing titer (NT50) using the formula: NT50=10^-IC50^. The neutralizing capacity of convalescent and vaccinee sera was measured against B.1 (D614G strain) and Omicron sublineages BA.1, BA.2, BA.4 and BA.5. For BTI sera, the neutralizing ability of sera was also assessed against the infecting variant (i.e. Gamma BTI were evaluated against Gamma and Delta BTI against Delta). To ensure equivalent infection levels, a ‘back-titration’ was performed in each experiment with each of the viral strains. Briefly, the viral dilution used to infect cells in the presence or absence of serum dilutions was titrated as above, in 10-fold dilutions in VGM and the TCID50 was calculated using the Reed and Muench formula to verify that the virus inoculum was 100 TCID50.

### Pseudotype preparation

HIV-based pseudotypes were generated as in (80) from the Firefly Luciferase-tagged HIV-1Δ*env*Δ*nef* backbone (81) complemented in trans with B.1, Omicron BA.1 or Omicron BA.2 Spike expression vectors lacking the 19 C-terminal amino acids containing the ER-retention signal (InvivoGen plv-spike-v11 and plv-spike-v12). HEK293T cells (1.2×10^6^ cells) were transfected with 2.5 μg HIV-1Δ*env*Δ*nef* and 0.5 (BA.1 and BA.2) or 1 μg (B.1) Spike expression vectors using JetPRIME. After 16 hours, medium was replaced and supernatants were collected after 48 hours, centrifuged for 5 minutes at 4°C and immediately used for neutralization assays.

### Pseudotype-based neutralization Assay

Serial three-fold serum dilutions starting at 1:40 were incubated 30 minutes at 4°C with B.1, Omicron BA.1 or Omicron BA.2 Spike pseudotypes. The serum/pseudotype mixture was then added to HEK293T-ACE2-TMPRSS2 cells (5×10^4^ cells/well) in DMEM containing 10% FBS for 60 hours. Cells were then lysed using the Promega lysis buffer (E1531 from Promega, Belgium) and a freeze-thay cycle and Firefly Luciferase activity was assessed using the Firefly Luciferase Assay following the manufacturer’s recommendations (E4550 from Promega, Belgium). Percent infection was calculated relative to infected cells in the absence of test serum. The IC50 was determined by inferring the 4-parameter non linear regression curve fit with unconstrained top and bottom values using GraphPad Prism v5 and was log-transformed into 50% neutralizing titer (NT50) as above.

### Statistical analyses

Statistical analyses were performed using GraphPad Prism v5. The Shapiro-Wilk test was used to verify distribution and all datasets were non-normally distributed. Differences between groups were compared using a Wilcoxon signed ranked test for comparisons between two groups and a Kruskal-Wallis signed-rank test followed by a Dunn’s post-hoc test for comparisons of three or more groups. A matched comparison was not applied in this case because data for all samples were not always available due to serum availability. Correlation coefficients (r) were determined using as Spearman’s rank correlation. P-values < 0.05 were considered significant.

## Results

### Early pandemic convalescent sera poorly neutralize Omicron strains

The convalescent sera used in this study were collected during the first SARS-CoV-2 wave from patients infected with the B.1 (Spike D614G) strain. All sera were collected during acute infection (median 16.7 days after symptom onset, interquartile range [IQR] = 13.5-19.9) and most were from patients with moderate or severe/critical COVID-19. Fifty percent neutralization titers (NT50) Geometric Mean (GMT) were comparable between B.1 (GMT=125.0, 95% Confidence interval (CI) [82.4, 189.7]) and Delta (GMT=153.3, 95% CI [94.36, 249.0]) (Fig 1A and 1B). At the highest serum concentration tested (1:40), a similar proportion of sera (p>0.05) failed to neutralize B.1 (19/58, 32.7%) and Delta (22/57, 38.6%) (Fig 1A). In contrast, all Omicron variants escaped neutralization by convalescent sera to some extent (Fig 1A, Fig 1C-1F and Supplementary Fig 1A-1D). BA.1 was the most resistant to neutralization by convalescent sera (neutralizing GMT=21.0 95% CI [19.3, 22.9]), with only one serum showing low-level neutralization, while all other sera were unable to even slightly neutralize Omicron BA.1 (Fig 1A and 1C). The NT50 of convalescent sera against BA.2 (GMT = 54.8, 95% CI [40.3, 74.5]), BA.4 (GMT = 37.2, 95% CI [26.6, 52.0]) and BA.5 (GMT = 60.4, 95% CI [40.8, 89.9]) were also significantly lower than against B.1 (Fig 1D-1F) and Delta (Supplementary Fig 1B-1D), but higher than BA.1 (Fig 1G-1I). BA.2 and BA.5 had similar sensitivities to neutralization with over 50% of non-neutralizing sera (BA.2: 27/52, 52% and BA.5: 25/47, 53.2%). BA.4 was slightly but significantly more resistant than BA.2 and BA.5 (p< 0.01 in both cases) and 75% (27/36) of the sera failed to neutralize this sublineage (Fig 1A, Fig 1C-1L). The full escape from neutralization of BA.1 and the relative residual sensitivity of BA.2 to neutralization were confirmed using HIV-1-SARS-CoV-2 Spike pseudotypes on a subset of sera (Fig 1M).

**Figure 1.**
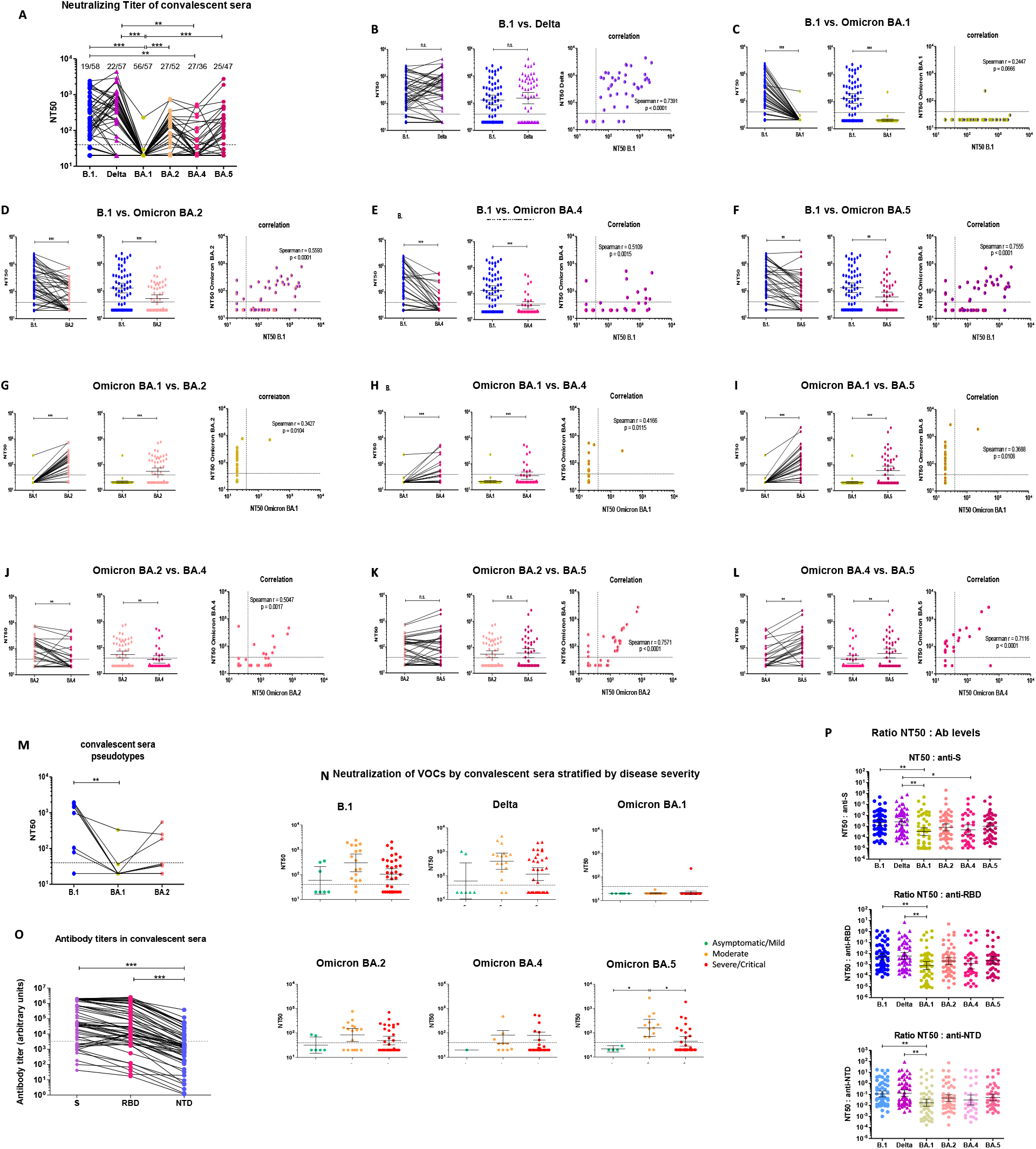
Neutralizing activity of pre-VOC unvaccinated SARS-CoV-2-infected convalescent sera against ancestral B.1, Delta and Omicron BA.1, BA.2, BA.4 and BA.5. A. Comparison of NT50 between all strains. The infecting strain is indicated on the x-axis and with color codes: blue circles = B.1, purple triangles = Delta, gold hexagons = Omicron BA.1, pink hexagons = Omicron BA.2, orange hexagons = Omicron BA.4 and burgundy hexagons = Omicron BA.5. This color code is used throughout the figure and manuscript. The dotted line represents the 1:40 serum dilution cut-off. The proportion of non-neutralizing sera is indicated above each data set. **B-L. Pairwise comparison and correlation of 50% neutralizing titers (NT50) between the tested strains**. Serial two-fold dilutions (starting 1:40, marked as a dotted line on graphs) of heat-inactivated convalescent sera were incubated with 100 TCID50 of the indicated viral strain. Vero-E6 cells (10^4^ cells/ well) were infected with the virus/serum mixture for 72 hours at 37°C and virus-induced cytophathic effect (CPE) was measured using a colorimetric assay based on the reduction of the tetrazolium salt WST-8 by live cells. All infections were performed in triplicate wells. Percent survival was calculated relative to uninfected cells (100% survival). The half-maximal inhibitory concentration (IC50) was determined by inferring the 4-parameter non linear regression curve fit (GraphPad Prism v5) with unconstrained top (100% survival) and bottom (no serum) values and log-transformed to assess the 50% neutralizing titer (NT50). Paired NT50 (left panel) and geometric mean titers (GMT) with 95% confidence intervals (95% CI) (middle panel) are shown. The correlation is shown on the right panel and the Spearman correlation coefficient (r) is indicated in the right panel for each pair. **M. NT50 for a subset of sera assessed using Luciferase-tagged HIV-1-Δ*env*Δ*nef*-Spike pseudotypes.** Three-fold serum dilutions starting at 1:40 were incubated with pseudotypes for 30 minutes, and then used to infect HEK293T-ACE2-TMPRSS2 cells (5×10^4^ cells/well). Firefly Luciferase activity was measured in cell lysates 60 hours after infection. Percent infection was calculated relative to infected cells in the absence of serum (100% infection), and the IC50 and NT50 were calculated as above. All infections were performed in triplicate. **N. NT50 of convalescent sera from patients with mild/asymptomatic (green), moderate (orange) or severe/critical (red) forms of COVID-19 against B.1, Delta, Omicron BA.1, BA.2, BA.4 and BA.5. O. Anti-Spike, anti-RBD and anti-NTD antibody levels in convalescent sera.** Antibody levels against Spike (purple), the Receptor Binding Domain (RBD) (pink) or the N-terminal domain (NTD) (blue) of Spike were measured in convalescent sera using the MSD V-plex platform for SARS-CoV-2. Antibody levels are reported as arbitrary units. **P. Ratios of NT50 to anti-S, (upper panel), anti-RBD (middle panel) and anti-NTD (lower panel) antibody levels.** For all analyses, differences between groups were compared using a Wilcoxon signed rank test for comparison between two groups (panels B-L) and a Kruskal-Wallis test followed by a Dunn’s multiple comparison post-hoc test for comparisons between three or more groups (panels A, M-P). P-values < 0.05 were considered significant. *: p<0.05; **: p<0.01; ***: p<0.001.

It is noteworthy that there was no continuum in the cross-neutralizing ability of convalescent sera against different strains. For instance, some sera failed to neutralize B.1 but neutralized Delta (Fig 1B) or even Omicron BA.2, BA.4 or BA.5 (Fig 1D-1F). More interestingly, some sera neutralized BA.5 better than BA.2 (Fig 1K). Most convalescent sera which cross-neutralized Omicron BA.2, BA.4, BA.5 had high NT50 (>350) against B.1 (Fig 1D-1F). Accordingly, there was a good, although imperfect, correlation between neutralizing activities of convalescent sera against B.1 and Delta (Spearman’s r = 0.7 391, p<0.0001) (Fig 1B) and a more modest correlation between B.1 or Delta and Omicron sublineages (Spearman r <0.6) (Fig 1C-1L and Supplementary Fig 1). Patients with moderate disease generally had higher neutralizing NT50 GMT than patients with mild or severe disease against all tested strains, although statistical support was reached only for BA.5 (Fig 1N).

To gain some qualitative insight on the antibodies mediating neutralization, we calculated the NT50:anti-S, NT50:anti-RBD and NT50:anti-NTD (N-terminal domain of S) ratios for all strains. This ratio subdivides the measured NT50 into the average neutralizing ability of individual antibodies and can thus be used as a surrogate to estimate antibody avidity (40, 46). As shown in Fig 1O, nearly all patients had detectable antibodies against S, the RBD and the NTD, although antibody levels varied substantially between patients. The NT50:anti-S, NT50:anti-RBD and N50:anti-NTD ratios were comparable for B.1 and Delta and were ~1 log_10_ lower for Omicron BA.1 (p<0.01) (Fig 1P). The three ratios were also lower for BA.2, BA.4 and BA.5, in line with the corresponding neutralizing titers. This observation suggests that antibodies elicited by infection with early SARS-CoV-2 partially cross-react with Omicron BA.2 and its sublineages BA.4 and BA.5 better than with BA.1.

### Sera from boosted vaccinees retain partial neutralizing activity against all Omicron sublineages

Next we assessed the neutralizing ability of sera from 14 triple-vaccinated individuals against the same SARS-CoV-2 VOCs. The time elapsed between booster dose and sampling varied between 15 days and 6 months (median = 4 months, IQR = 2.26 – 6). Infection history is not known. The NT50 GMT for B.1 and Delta were 246.8, 95% CI [124.4, 489.8] and 344.6, 95% CI [155.6, 763.3]) respectively. The correlation between NT50s of both pre-Omicron VOCs was very good (Spearman r=0.9163, p<0.0001), indicating that sera which neutralized B.1 also neutralized Delta (Fig 2A and 2B). As shown in Fig 2A and 2B, 2/14 vaccinee sera (14%) were unable to neutralize the ancestral B.1 and Delta. Time since vaccination for these two samples was 5 and 6 months. There was a significant drop in neutralization of all Omicron lineages compared to B.1 and Delta (Fig 2C-2F and Supplementary Fig 2A-2D). The NT50 GMTs were lowest for BA.1 (NT50 = 51.1 (95% CI [25.6, 100.5]), BA.4 (NT50 = 46.1, 95% CI [25.7, 82.7]) and BA.5 (NT50 = 58.9 (95% CI [34.9, 99.5] (Fig 2C-2L). Despite a significantly lower NT50 GMT compared to B.1 (Fig 2D), BA.2 remained significantly more sensitive to neutralization by vaccinee sera than the other Omicron sublineages (NT50 = 119.2 (95% CI [55.5, 255.8]) (Fig 2G, Fig 2J and Fig 2K), with only 2 non-neutralizing sera at the 1:40 dilution, while a higher proportion of sera failed to neutralize the other Omicron sublineages at the highest dilution tested (7/14 for BA.1, 8/14 for BA.4 and 5/14 for BA.5) (Fig 2A). Overall, in contrast to convalescent sera, the neutralizing ability of vaccinee sera against B.1 extended to other VOCs, i.e. sera which poorly neutralized B.1 generally failed to neutralize Omicron sublineages and neutralizing sera retained some neutralizing ability against Omicron strains as well (Fig 2A, Fig 2G). Accordingly, the side-by-side comparison of sera NT50 against pre-Omicron (B.1 and Delta) and Omicron VOCs showed overall good Spearman correlation coefficients (Spearman r > 0.66) (Fig 2C-2L and Supplementary Fig 2A-2D). Similar results for BA.1 and BA.2 were again recorded with pseudotypes (Fig 2M).

**Figure 2.**
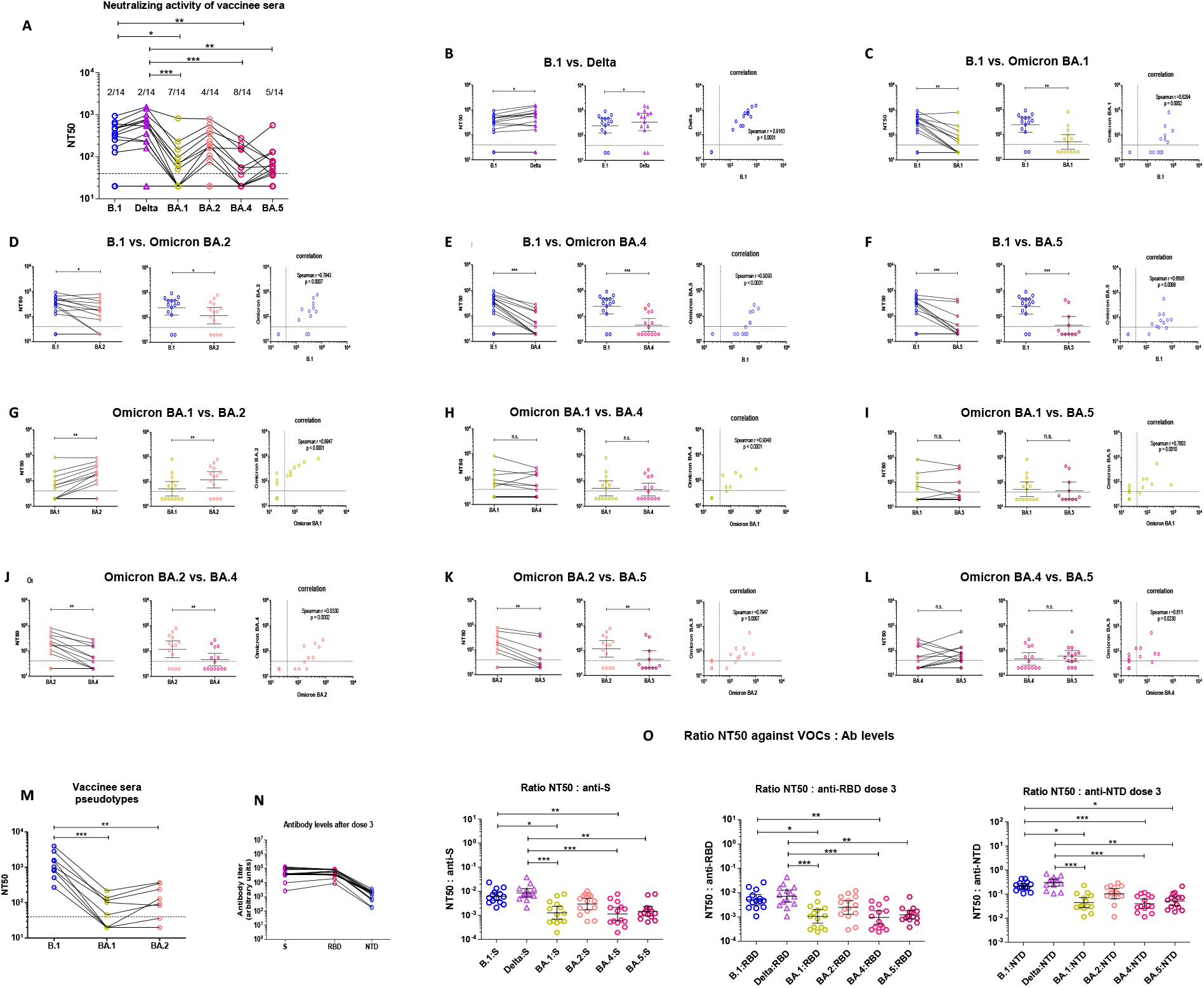
Neutralizing activity of sera from triple-vaccinated individuals against ancestral B.1, Delta and Omicron BA.1, BA.2, BA.4 and BA.5. A. Comparison of NT50 between all strains. The infecting strain is indicated on the x-axis and with color codes: blue open circles = B.1, purple open triangles = Delta, gold open hexagons = Omicron BA.1, pink open hexagons = Omicron BA.2, orange open hexagons = Omicron BA.4 and burgundy open hexagons = Omicron BA.5. This color code is used throughout the figure and manuscript. The dotted line represents the 1:40 serum dilution cut-off. The proportion of non-neutralizing sera is indicated above each data set. **B-L**. **Pairwise comparison and correlation of 50% neutralizing titers (NT50) between the tested strains.** Serial two-fold dilutions (starting 1:40, marked as a dotted line on graphs) of heat-inactivated sera from vaccinees 3.45 months, 95%CI [2.26, 5.25] after dose 3 were incubated with 100 TCID50 of the indicated viral strain. Vero-E6 cells (10^4^ cells/ well) were infected with the virus/serum mixture for 72 hours at 37°C and virus-induced cytophathic effect (CPE) was measured using a WST-8-based colorimetric assay. All infections were performed in triplicate wells. Percent survival was calculated relative to uninfected cells (100% survival). The IC50 and NT50 were calculated. Paired NT50 (left panel) and geometric mean titers (GMT) with 95% confidence intervals (95% CI) (middle panel) are shown. The Spearman correlation coefficient (r) is indicated in the right panel for each pair. **M. NT50 for a subset of sera assessed using Luciferase-tagged HIV-1-Δ*env*Δ*nef*-Spike pseudotypes.** Three-fold serum dilutions starting at 1:40 were incubated with pseudotypes for 30 minutes, and then used to infect HEK293T-ACE2-TMPRSS2 cells (5×10^4^ cells/well). Firefly Luciferase activity was measured in cell lysates 60 hours after infection. Percent infection was calculated relative to infected cells in the absence of serum (100% infection), and the IC50 and NT50 were calculated as above. All infections were performed in triplicate. **N. Anti-Spike, anti-RBD and anti-NTD antibody levels in vaccinee sera.** Antibody levels against Spike (purple), the Receptor Binding Domain (RBD) (pink) or the N-terminal domain (NTD) (blue) of Spike were measured in vaccinee sera using the MSD V-plex platform for SARS-CoV-2. Antibody levels are reported as arbitrary units. **O. Ratios of NT50 to anti-S, (left panel), anti-RBD (middle panel) and anti-NTD (right panel) antibody levels.** For all analyses, differences between groups were compared using a Wilcoxon signed rank test for comparison between two groups (panels B-L) and a Friedman test followed by a Dunn’s multiple comparison post-hoc test for comparisons between three or more groups (panels A and M-O). P-values < 0.05 were considered significant. *: p<0.05; **: p<0.01; ***: p<0.001.

Next we calculated the NT50:antibody ratios for vaccinees. All vaccinees had detectable antibodies against S, the RBD and the NTD in serum (Fig 2N). Overall, antibody levels spanned a narrower range than convalescent sera and most antibodies against S target the RBD. Again, the NT50:antibody level ratio was comparable for B.1 and Delta, but was markedly lower (~1 log_10_) for Omicron BA.1, BA.4 and BA.5 for all antibodies (anti-S, anti-RBD and anti-NTD) (Fig 2N), illustrating the lower affinity of vaccine-induced antibodies for these Omicron sublineages. Despite identical Spike sequences, the NT50:Ab ratio was lower for BA.4 than for BA.5, reflecting subtle differences in the susceptibility of these two sublineages to neutralization. The NT50:antibody ratios for Omicron BA.2 were intermediate, nicely recapitulating the NT50 profiles. These figures indicate that antibodies elicited by first generation vaccines retain sufficiently high affinity for BA.2 Spike determinants, whilst BA.1, BA.4 and BA.5 have evolved to further escape binding and thereby neutralization by pre-Omicron-elicited antibodies.

Taken together, these results document that antibodies elicited by first generation vaccines partially retain neutralizing ability against Omicron sublineages 4 months after the 3^rd^ dose, despite a significant drop compared to the ancestral B.1. They also clearly show differences between infection-elicited and vaccine-elicited antibodies: convalescent sera were unable to neutralize Omicron BA.1 but they similarly neutralized Omicron BA.2, BA.4 and BA.5, whereas vaccinee sera partly neutralized Omicron BA.2 but had similarly low neutralizing ability against Omicron BA.1, BA.4 and BA.5 (compare Fig 1A and Fig 2A).

### Breakthrough infection sera have distinct neutralization profiles and retain cross-neutralizing ability against all Omicron sublineages

Hybrid immunity conferred by vaccination and infection together was reported to be superior to immunity elicited by infection or vaccination alone (3, 20, 37, 38, 65, 66, 70, 73, 76, 82–85). We therefore further investigated the susceptibility of the four Omicron sublineages to sera from 16 vaccinated individuals with breakthrough infection (BTI) with pre-Omicron VOCs (2 infected with Gamma and 14 with Delta). BTI patients were infected between July 15^th^ and September 20^th^ 2021, when Gamma and Delta were the main circulating VOCs in Luxembourg. The median time elapsed since the 2^nd^ vaccine dose was 3.1 months [CI=2.1-4.5]. Most sera were collected at the time of diagnosis but time since symptom onset is unknown. As shown in Fig 3A and 3B, half the sera were strongly neutralizing against B.1 and the infecting VOC (Gamma or Delta), while the other half was fully non-neutralizing (5 sera: 4 Delta, 1 Gamma) or poorly neutralizing (3 sera). Overall NT50 was not significantly different between B.1 and the infecting VOC (Gamma for Gamma-BTI cases and Delta for Delta-BTI cases): GMT B.1 = 194.6, 95% CI [65.6, 577.0]) and GMT Gamma/Delta =329.5, 95% CI [102.0, 1064.0], p<0.05). All Omicron lineages escaped neutralization by BTI sera to some extent (Fig 3A and Fig 3C-3F), with the following NT50 GMT: BA.1 NT50=44.4, 95% CI [25.0, 78.9], BA.2 NT50=70.7, 95% CI [32.3, 154.9], BA.4 NT50=51.6, 95% CI [25.4, 104.9], and BA.5 NT50= 87.9, 95% CI [34.4, 224.6]) (Fig 3C-3F and Supplementary Fig 3A-3D). As for convalescent and vaccinee sera, the drop in neutralization was more pronounced for BA.1, although this trend did not reach statistical significance, probably due to the low number of cases and the high proportion of non-neutralizing BTI sera (Fig3A and Fig 3G-3L). Strikingly, Delta BTI sera retained good neutralizing ability against BA.5, as illustrated by the excellent correlation score (Fig 3F). BA.4 was slightly more resistant to neutralization than BA.2 and BA.5 (p<0.05 in both cases) (Fig 3J and 3L), as previously recorded for unvaccinated convalescent sera.

**Figure 3.**
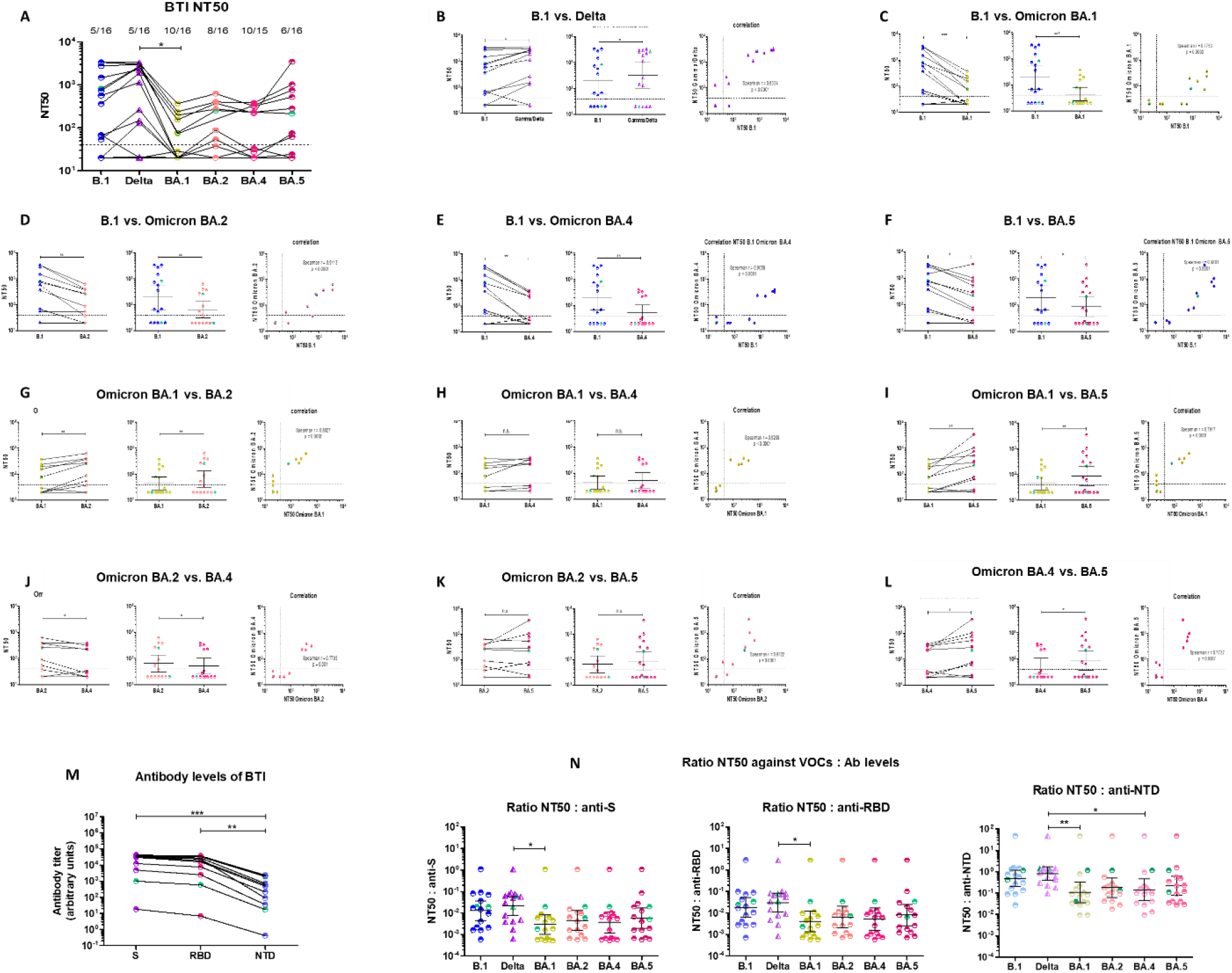
Neutralizing activity against ancestral B.1, Delta and Omicron BA.1, BA.2, BA.4 and BA.5 of sera from vaccinated individuals (2 doses) with Delta or Gamma breakthrough infection (BTI). A. Comparison of NT50 between all strains. The infecting strain is indicated on the x-axis and with color codes: partially filled blue circles = B.1, partially filled purple triangles = Delta, partially filled gold hexagons = Omicron BA. 1, partially filled pink hexagons = Omicron BA.2, partially filled orange hexagons = Omicron BA.4 and partially filled burgundy open hexagons = Omicron BA.5. Gamma-BTI are identified with green symbols in all panels. The dotted line represents the 1:40 serum dilution cut-off. This color code is used throughout the figure. The proportion of non-neutralizing sera is indicated above each data set. **B-L**. **Pairwise comparison and correlation of 50% neutralizing titers (NT50) between the tested strains.** Serial two-fold dilutions (starting 1:40, marked as a dotted line on graphs) of heat-inactivated sera from BTI patients collected at the time of diagnosis were incubated with 100 TCID50 of the indicated viral strain. Vero-E6 cells (10^4^ cells/ well) were infected with the virus/serum mixture for 72 hours at 37°C and virus-induced cytophathic effect (CPE) was measured using a WST-8-based colorimetric assay. All infections were performed in triplicate wells. Percent survival was calculated relative to uninfected cells (100% survival). The IC50 and NT50 were calculated. Paired NT50 (left panel) and geometric mean titers (GMT) with 95% confidence intervals (95% CI) (middle panel) are shown. The Spearman correlation coefficient (r) is indicated in the right panel for each pair. **M. Anti-Spike, anti-RBD and anti-NTD antibody levels in BTI sera.** Antibody levels against Spike (purple), the Receptor Binding Domain (RBD) (pink) or the N-terminal domain (NTD) (blue) of Spike were measured in vaccinee sera using the MSD V-plex platform for SARS-CoV-2. Antibody levels are reported as arbitrary units. **O. Ratios of NT50 to anti-S, (left panel), anti-RBD (middle panel) and anti-NTD (right panel) antibody levels.** For all analyses, differences between groups were compared using a Wilcoxon signed rank test for comparison between two groups (panels B-L) and a Kruskal-Wallis test followed by a Dunn’s multiple comparison post-hoc test for comparisons between three or more groups (panels A, M and N). P-values < 0.05 were considered significant. *: p<0.05; **: p<0.01; ***: p<0.001.

Similarly to the cross-neutralizing profile of vaccinee sera, the cross-neutralizing profile of BTI sera was also relatively well maintained across strains. Sera which neutralized B.1 at dilutions higher than 1:80 had similar or higher neutralizing ability against the corresponding infecting VOC (Fig 3B) and six retained some, although significantly weaker, cross-neutralization against all Omicron sublineages. Conversely, most sera which did not neutralize B.1 also failed to neutralize the infecting VOC (1 Gamma and 3 of 4 non-neutralizing Delta-BTI sera) (Fig 3B) and Omicron sublineages (Fig 3C-3L and Supplementary Fig 3A-3D). Accordingly, there was a very good correlation between the NT50s of BTI sera against B.1, the infecting VOC (Gamma or Delta) (Spearman r = 0.9334, p<0001) as well as against the Omicron sublineages (Spearman r> 0.77 in all) (Fig 3C-3L and Supplementary Fig 3A-3D).

Despite distinct neutralization profiles, all but one BTI sera had antibodies against S, the RBD and the NTD (Fig 3M). The NT50:antibody ratios were again ~1 log_10_ higher for B.1 and Delta compared to Omicron strains, although statistical support was reached only between the infecting strain and BA.1 for the three ratios and for the NT50:anti-NTD ratio as well for BA.4 (Fig 3N).

Overall, BTI sera showed a distinct neutralization profile, with two groups of sera, neutralizing or non-neutralizing. There was a good cross-neutralization across strains, including Omicron, as recorded for vaccinee sera and a small loss in antibody avidity estimated from the NT50:antibody level ratio.

### Comparison of convalescent, vaccinee and BTI sera

Because there were shared trends and differences between the three groups of sera included in this study, we further compared the NT50s and NT50:antibody ratios of all pooled sera on one hand, and the three groups side by side on the other.

The comparison of the NT50s of all sera (convalescent + vaccinee + BTI) highlighted a trend to higher neutralization of Delta compared to B.1, and a clear drop (~1.5 log_10_) in neutralizing ability against all Omicron sublineages compared to B.1 and Delta (Fig 4A). The strongest escape from neutralization was recorded for Omicron BA.1, the weakest for BA.2 and BA.5, while BA.4 was intermediate (Fig 4A). Antibody avidity, inferred from the NT50:antibody ratios, was significantly lower for all Omicron sublineages in comparison to B.1 and Delta (Fig 1B). Again the most spectacular drop was recorded for BA.1 and BA.4, while BA.2 and BA.5 had comparable NT50:antibody ratios for S, the RBD and the NTD. Vaccinee and BTI sera had the highest NT50s (Fig 4A) and the highest ratios were among convalescent and BTI sera (Fig 4B).

**Figure 4.**
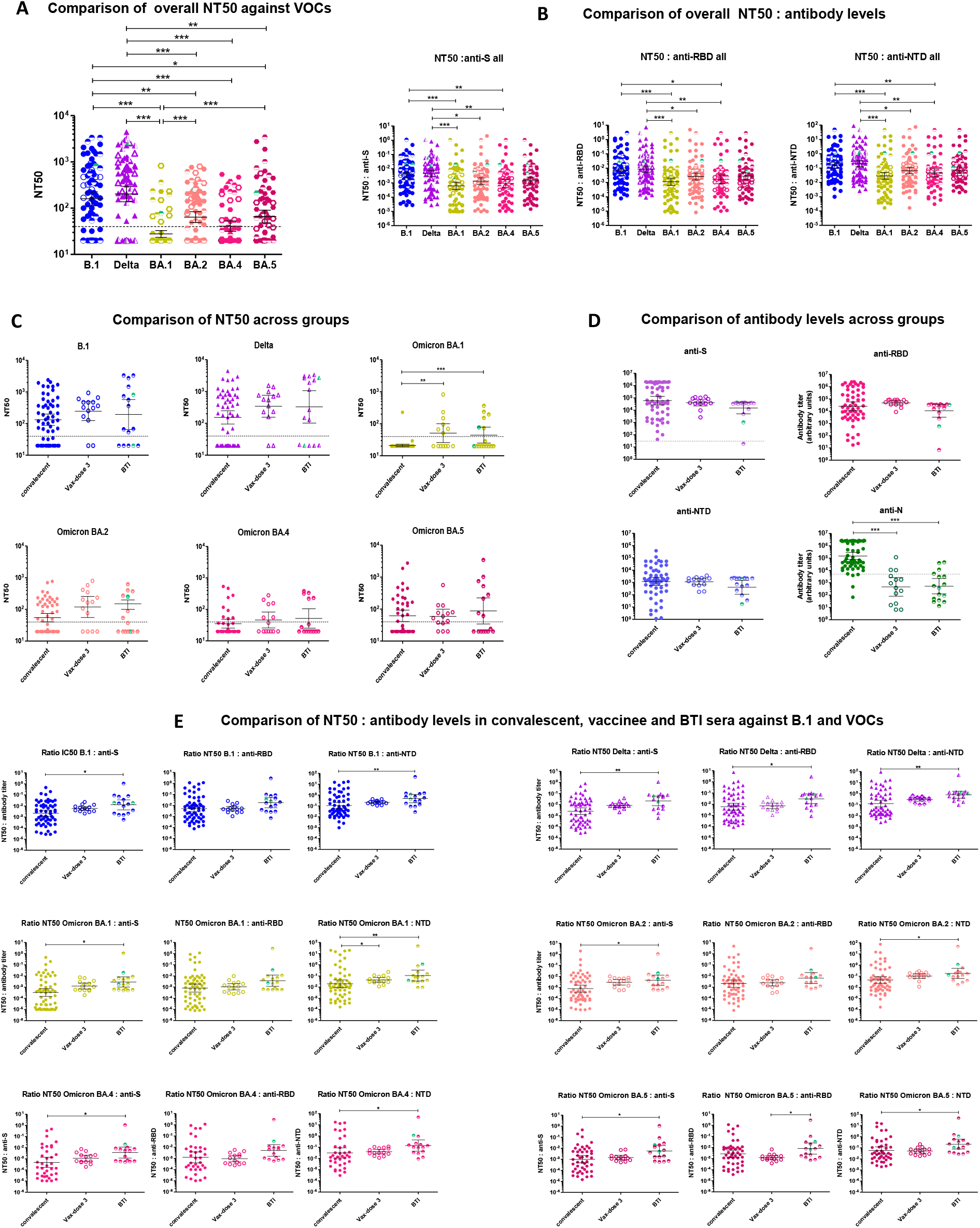
Comparison of neutralizing activities of convalescent, vaccinee and BTI sera against B.1, Delta and Omicron BA.1, BA.2, BA.4 and BA.5. A. Overall comparison of NT50 of all sera (convalescent + vaccinee + BTI) against B.1, Delta and Omicron BA.1, BA.2, BA.4 and BA.5. NT50 from Fig 1, Fig 2 and Fig 3 were analyzed together and compared. The dotted line represents the 1:40 serum dilution cut-off. **B. Overall comparison of NT50:antibody ratios of all sera (convalescent + vaccinee + BTI).** NT50:anti-S, NT50:anti-RBD and NT50:NTD ratios from Fig 1, Fig 2 and Fig 3 were analyzed together and compared**. C. Comparison of NT50s from convalescent, vaccinee and BTI sera for B.1, Delta, Omicron BA.1, BA.2, BA.4 and BA.5. NT50 of convalescent, vaccinee and BTI sera were compared for each strain.** The infecting strain is indicated above each panel. For BTI sera, the Gamma-BTI patients infected with Gamma are represented with green symbols. **D. Comparison of antibodies against the Nucleocapsid (N), Spike (S), RBD and NTD in convalescent, vaccinee and BTI sera.** For BTI sera, the Gamma-BTI are represented with black circles and the Delta-BTI are represented in purple for anti-S antibodies, pink for anti-RBD antibodies, blue for anti-NTD antibodies and green for anti-N antibodies. Anti-N and anti-S antibodies were also estimated with the Eurimmun assay for BTI and only 3 BTI sera had anti-N antibodies > 0. The grey dotted lines in the panels for anti-S and anti-N antibody levels mark the threshold between anti-S and anti-N-positive and negative sera based on the Eurimmun assay. **E. Comparison of the NT50:antibody level ratios for convalescent, vaccinee and BTI sera.** The NT50:anti-S, NT50:anti-RBD and NT50:NTD ratios for convalescent, vaccinee and BTI sera against each strain are compared. The infecting strain is indicated above each group of panels. For BTI sera, the Gamma-BTI are represented with green symbols. For all analyses, differences between groups were compared using a Kruskal-Wallis test followed by a Dunn’s multiple comparison post-hoc test. P-values < 0.05 were considered significant. *: p<0.05; **: p<0.01; ***: p<0.001.

We then compared the NT50s from convalescent, vaccinee and BTI sera. Overall, convalescent sera had the lowest NT50 GMT and BTI sera had the highest NT50 GMT (Fig 4C). This trend held true for all strains, although statistical support was reached only for Omicron BA.1, reflecting the fact that BA.1 fully escaped neutralization by convalescent sera. Vaccinee and BTI sera had similar NT50 GMT for most strains, but when the non-neutralizing BTI sera were excluded from the analysis, BTI sera featured significantly higher NT50s (p>0.001) (Supplementary Fig 4). Of note, neutralization of BA.4 and BA.5 by convalescent, vaccinee and BTI sera was comparable, reflecting the better cross-neutralization of these sublineages by convalescent sera (Fig 1A), and the poor neutralizing ability of vaccinee sera against BA.4 (Fig 2A). It is worth mentioning that convalescent sera were collected at the time of acute infection, while 4 months had elapsed since the last vaccine dose for vaccinees and 3 months for BTI. Therefore, neutralizing ability 4 months after the 3^rd^ vaccine dose remained higher than that conferred by early infection against most strains tested (B.1, Delta, Omicron BA.1 and BA.2) (Fig 4C).

Although convalescent sera had lower NT50s, the levels of antibodies targeting Spike and the NTD were comparable between convalescent and vaccinee sera (Fig4D). BTI sera tended to have overall lower antibody levels compared to convalescent and vaccinee sera (Fig 4D). Anti-RBD antibodies tended to be higher in vaccinee sera compared to convalescent and BTI sera (Fig 4D), probably reflecting the open conformation of the Spike in mRNA vaccines. Given the time elapsed since the third vaccine dose, these figures likely underestimate these qualitative differences. Therefore similar antibody levels ensured higher neutralization in vaccinated individuals (both uninfected and BTI) than in acutely infected unvaccinated patients. Although the time since symptom onset is not known, the levels of anti-N antibodies (Fig 4D) confirm that in most cases, infection of BTI was recent, and probably had triggered a rapid boost of anti-S, anti-RBD and anti-NTD antibodies from memory B-cells, but not yet the appearance of anti-N antibodies.

The NT50:anti-S and NT50:anti-NTD ratios in BTI sera were higher than those in convalescent and vaccinee sera (Fig 4E). The difference was statistically significant between BTI and convalescent sera for all strains. Again, these differences are likely underestimated, given that half the BTI have no neutralizing ability. The NT50:anti-S and NT50:anti-NTD ratios of vaccinee sera were intermediate between convalescent sera and BTI (Fig 4E). Anti-RBD antibodies showed a similar trend between convalescent and BTI sera, but statistical support was reached only for Gamma/Delta, i.e., for the infecting strain. Importantly, the NT50:RBD ratio for vaccinee sera was similar to convalescent sera, indicating that the slightly higher anti-RBD levels likely account for the higher NT50, suggesting a more targeted immune response.

Together, these results indicate that breakthrough infection elicits neutralizing antibody responses with higher avidity than infection alone, but also than booster vaccination, including against Omicron sublineages. They also indicate that the higher cross-neutralizing ability of BTI sera is likely due to the superior avidity of antibodies (Fig 4E). This may explain, at least partly, the superiority of hybrid immunity over infection or vaccination alone, and is consistent with ongoing affinity maturation after infection and vaccination(37, 40, 41, 82, 84, 86, 87).

## Discussion

In this study, we compared the neutralizing ability of pre-Omicron immune sera from convalescent, triple-vaccinated and pre-Omicron breakthrough infection individuals against four Omicron sublineages. All sera exhibited a marked or full drop in neutralizing ability against all Omicron sublineages compared to B.1 and Delta (Fig 4). BA.1 was the most resistant to neutralization and BA.2 the least (Fig 1A, Fig 2A and Fig 3A). BA.4 and BA.5 had intermediate resistance levels overall, albeit with differences between the two sublineages and between the groups of sera (Fig 4). Convalescent and BTI sera poorly neutralized BA.1, neutralized BA.4 slightly better, and similarly neutralized BA.2 and BA.5 (BA.1<BA.4<BA5~BA.2). Vaccinee sera, in contrast, neutralized Omicron BA.5 significantly less efficiently than BA.2 (Fig 2K) (BA.1<BA.4<BA.5<BA.2), in line with prior studies reporting the higher escape of BA.5 from NAbs (2, 4, 16, 30, 31, 50, 62–64, 88). The high evasion capacity of Omicron BA.1, BA.4 and BA.5 to immunity from vaccination with two, three or four doses is extensively documented (2–4, 16, 18, 20, 24, 30, 40, 43, 44, 47–55, 57, 59–66, 72). Our results however reveal that convalescent sera retain some neutralizing ability against the BA.5 sublineage. In contrast, they were unable to neutralize BA.1, in agreement with previous reports (42, 57, 75). This observation somehow mirrors the restricted cross-neutralizing ability of BA.1-elicited antibodies against pre-Omicron and other Omicron lineages (3, 20, 30, 42, 50, 63, 66, 73, 75–77, 89), and highlights that BA.1 is antigenically very distant from all other VOCs and triggers strongly imprinted immune responses. Accordingly one study found that BA.1 booster triggers the activation of naïve B-cells rather than memory B-cells (74), while others record a small proportion of naïve B-cell activation (2, 63, 67, 87). This observation also suggests that different antigens (infection or mRNA vaccines) lead to slightly different immune imprints. BA.4 and BA.5 share identical Spike sequences and differ by 3 mutations located in ORF7b (L11F in BA.4), in N (P151S in BA.4) and in the Membrane protein M (D3S in BA.5). The widespread use of pseudotypes does not allow to pick-up differences between BA.4 and BA.5 and only few reports have investigated the susceptibility of BA.4 to NAbs using live virus (3, 66). These mutational differences confer BA.5 a growth advantage over BA.4. This is consistent with the observation that BA.4 has not spread much in Luxembourg, and BA.5 has become the dominant variant in Luxembourg and elsewhere, indicating that infectivity and transmissibility rather than NAb escape were the main selective drivers, as previously reported (**4, 20**). Our findings however indicate that these mutations may also allow a better escape from NAbs through indirect mechanisms which would warrant to be explored further.

Despite differences between the neutralizing abilities of convalescent, BTI and vaccinee sera, all groups of sera showed some resilience towards BA.2 (Fig 1, Fig 2 and Fig3). The literature on the susceptibility of BA.2 to neutralization is still controversial, as some studies record similar or higher resistance to neutralization for BA.2 compared to BA.1 (4, 16, 48, 50, 61, 62, 85) while others record a less dramatic drop in neutralization for BA.2 compared to BA.1 (4, 20, 33, 43, 64, 73). These differences probably ensue from different vaccination/infection histories and from different experimental and calculation approaches (target cells, live virus versus pseudotypes, PRNT50/PRNT90, NT50). Our findings with both live virus and pseudotypes agree with the latter. Furthermore, the NT50:antibody ratios in vaccinee sera were higher for BA.2 compared to the other Omicron sublineages, indicative of residual affinity (Fig 2O). The fact that BA.2 remains partially susceptible to neutralization by convalescent and vaccinee sera while BA.4 and BA.5 more efficiently eluded humoral responses suggests that its selective advantage lied in increased transmissibility at a time where vaccine and infection coverage was still moderate. As increasing numbers of individuals acquired immunity through infection or vaccination, the relative advantage of infectivity over immune escape must have shifted. Accordingly, a number of BA.2-derived offspring are spurring in a BA.5-dominated context. The acquisition of supplementary mutations in Spike, such as R346K (BA.2.75.2) or F486P (BA.2.10.4 and BA.4.6) increases their ability to evade NAb and illustrates the strength of the immune selective pressure imposed on the virus (4, 30, 50, 64). While some of these mutations arise independently in different sublineages (converging evolution), others appear to be more specific (2). The increasing number of mutations accumulated by these variants underscore not only their role in incrementing antibody evasion and viral fitness, but also the plasticity of the SARS-CoV-2 genome. More importantly, they clearly indicate that SARS-CoV-2 has not reached an evolutionary threshold and that there is still room for evolution.

NAb levels are thought to be predictive of protection (90, 91) and are inversely correlated with viral load (85). In our study, convalescent sera and vaccinee sera harbored comparable anti-S, anti-RBD and anti-NTD antibody levels, while BTI tended to have slightly lower antibody levels (Fig 4D). These quantitative similarities likely reflect sampling times (acute infection for convalescent and BTI, ~4 months after 3^rd^ vaccine dose for vaccinees and ~3 months after 2^nd^ vaccine dose for BTI). However, there were profound qualitative differences in antibodies from unvaccinated convalescent sera compared to vaccinee sera, in agreement with other recent studies, albeit based on different BTI infections (3, 41, 42, 66). First, sera from vaccinated individuals (both uninfected and BTI) which neutralized B.1 cross-neutralized Delta (Fig 2A and 2B and Fig 3A and 3B) and partially neutralized Omicron sublineages, while convalescent sera had a less straightforward cross-neutralization pattern, as some non-neutralizing sera against B.1 were able to neutralize Delta or Omicron sublineages (Fig 1A-L). Second, antibodies in convalescent sera mediated lower neutralization against all strains than vaccinees and BTI, in agreement with previous studies by us and by others (3, 46, 57, 61, 66, 83, 85). This was particularly striking for Omicron BA.1, which all convalescent sera were unable to neutralize (Fig 1A). In contrast, vaccinee sera, with or without breakthrough infection, partially retained cross-neutralizing ability against all Omicron sublineages, including against the notoriously resistant BA.1, BA.4 and BA.5 (Fig 2 and Fig 3). Thus, similar antibody levels induced by vaccination mediated significantly different levels of cross-neutralization, indicative of superior efficacy and avidity, particularly in the case of hybrid immunity (BTI infections with neutralizing ability). In all groups of donors, the NT50:anti:RBD ratio of vaccinee sera was lower for BA.1, BA.4 and BA.5 (Fig 4E), reflecting a much lower avidity of antibodies targeting the RBD against these sublineages, as previously reported by others (3, 20, 77). Antibody avidity (estimated from the NT50:antibody ratios) slightly but not significantly lower in convalescent than in vaccinee sera, and was significantly higher in BTI sera (Fig 4B and 4E). This observation likely reflects the fact that convalescent sera were collected during acute infection, while there had been time for affinity maturation in vaccinee and BTI sera. It is however noteworthy that unvaccinated patients with moderate disease had higher neutralizing NT50 GMT than patients with mild or severe disease against all strains (Fig 1N), revealing that these patients mount a neutralizing response comparable to that triggered by vaccination. Together, our findings confirm the superiority of hybrid immunity (3, 20, 37, 38, 50, 65, 66, 70, 73, 76, 82–85), including that conferred by pre-Omicron antigens towards Omicron strains, in agreement with previous studies (3, 41, 42, 46, 57, 61, 66, 76, 83, 85). These findings held true for all strains, illustrating the qualitative superiority of hybrid immunity over infection or vaccination but was particularly striking for Delta, in line with the fact that BTI were infected with Delta. Accordingly, the NT50:anti-RBD ratio in BTI sera for Delta was significantly higher than in convalescent sera. This observation can reflect affinity maturation after vaccination and/or after infection or beneficial immune imprinting which results in very effective binding to the Delta RBD, although it is not possible to distinguish between the two processes from our data. Nevertheless, since our study only included pre-Omicron sera, affinity maturation and immune imprinting between convalescent, vaccinated and BTI are expected to share multiple paratopes, with the exception of BTI sera, which share T478K and L452R with BA.4 and BA.5.

Converging evidence indicates that the time since vaccination/infection as well as the infecting variant dictate the level and quality of immune responses to mRNA boosters vaccination (70, 79). Ongoing affinity maturation shapes immune responses elicited by infection (82, 92–95) and by vaccine boosters, including those based on the original Wuhan strain (37, 40, 41, 82, 84, 86, 87). Accordingly, the time elapsed between vaccination and breakthrough infection is proportional to the breadth of the resulting antibody response (84). We do not have information on the time window between the second and the third vaccine doses in vaccinees included in this study, but based on the typical vaccination protocols deployed in Luxembourg and in Europe at the time, it is most likely that it exceeded 6 months, thus longer than the 4 months that elapsed between the boost and breakthrough infection in the BTI group. However, we found no correlation between the time window between vaccination (3^rd^ dose) and breakthrough infection and NT50 against the infecting strain (not shown) but this may be due to the low number of BTI cases.

With the burst in breakthrough infections due to Omicron, numerous studies have investigated cross-neutralization between Omicron-induced NAbs and other Omicron sublineages. Converging evidence documents the narrow antibody profile and the strong immune imprinting conferred by BA.1 infection (3, 20, 30, 42, 50, 57, 63, 66, 76, 85), suggesting it drove the current BA.2-derived Omicron landscape (e.g. R364K and L542 substitutions) (2, 63, 67). BA.2 imprinting seems to be less restrictive (2, 4, 16, 30, 31, 64, 77). Likewise, the numerous reports on ongoing affinity maturation after pre-Omicron immunity triggered by infection (82, 92–95) and by vaccination (37, 40, 41, 66, 82, 84, 86) suggest a broader immune window. In terms of unraveling cross-protection, however, the picture is blurred by the combination of data from vaccinated (hybrid immunity including a “Wuhan background”), re-infected individuals and unvaccinated individuals (“pure Omicron immunity). It is therefore difficult to discern the contribution of cross-neutralization due to shared epitopes and affinity maturation of a Wuhan-elicited immunity from cross-neutralization due to shared epitopes between Omicron sublineages. This observation is of particular importance in light of the deployment of second-generation vaccines, and on the open questions raised by the risk of obstructive immune imprinting. As infections with different strains and VOCs continue to swarm, a plethora of different immune profiles are emerging, reflecting infections with different VOCs, before or after vaccination, distinct vaccination schemes … The use of Wuhan-based or of second generation bivalent vaccines including different Omicron sublineages will add a further layer of complexity to the individual immune repertoires. Therefore, at a population level, the dynamics of the pandemic assemble into a puzzle of immune profiles. This collective immune evolution represents a strong evolutionary pressure for circulating strains. As of now, large scale immunization campaigns based on similar viral (Wuhan strain) sequences, and the global spread of VOCs in successive albeit rather homogeneous waves, have shaped immune responses in a globally similar fashion, despite some geographic specificities (Gamma in Brazil, Epsilon or Iota in the US…). The burst in Omicron infections and breakthrough infections has accelerated the impact of immune imprinting, causing significant reduction in NAbs. This concentrated humoral immune pressure in turn promotes convergent evolution (2, 63, 67, 77) and suggests the SARS-CoV-2 landscape may evolve towards the coexistence of a constellation of different strains with shared immune evasion and adaptive mutation patterns. The spurring of new variants derived from BA.2. (BA.2.12.1, BA.2.75, BA.2.75.2, BA.2.10.4…), from BA.4 (BA.4.6) or from BA.5 (BQ.1.1.) with common immune escape mutations testifies of the reduction of NAb epitope diversity. The ability of multivalent vaccines to overcome the immune evasion potential conferred by these mutations and to trigger memory B-cell affinity maturation will be crucial in breaking this vicious cycle. Indeed, recent studies reported that BA.1 breakthrough infection not only induces broadly NAbs, but also reactivates cross-reactive memory B-cell responses which accumulate somatic mutations and increase the breadth of NAbs (67, 87), while another study indicates it induces novel naïve B-cells (74). These two mechanisms are not mutually exclusive and the resulting expansion of the memory B-cell repertoire is consistent with the superiority of hybrid immunity and the cross-reactivity of NAbs conferred by Omicron breakthrough infections. Importantly, this increased breadth and potency was not seen at the serological level but only at the memory B-cell level (67), suggesting immune imprinting should be evaluated beyond serology. In this scenario, pan-coronavirus and vaccines that specifically target conserved and T-cell epitopes may also play a major role. The two open questions in the evolving SARS-CoV-2 landscape are how the virus will evolve, i.e. whether there is still sufficient evolutionary room for the Omicron lineage to evolve further towards full escape from a broad range of NAbs without reaching the threshold of lethal mutagenesis, and whether immune imprinting may be overcome by immunization with antigenically distinct strains or vaccines through affinity maturation or through the activation of other naïve B-cells.

The data reported here only includes pre-Omicron sera and thus sheds light on pre-Omicron cross-immunity. It would have been insightful to compare the cross-neutralization and particularly antibody avidity in sera from pre-Omicron and Omicron breakthrough infections, in order to gain further insight on if and to what extent Delta and Omicron breakthrough infections mold immune responses. There is high vaccine coverage in Luxembourg and thus good protection against severe forms of COVID-19, resulting in less hospitalizations. Indeed, breakthrough infections and hospital admissions increased from Wuhan to Delta, but decreased overall with Omicron, despite increased infectivity of the later Omicron sublineages (19, 20). With the coming Fall and Winter and the ballet of novel BA.2 offspring, epidemiological surveillance coupled to rapid assessment of susceptibility to NAbs and T-cell responses will be essential to orient vaccination strategies. Another shortcoming of this study is that it only includes a small number of vaccinee and BTI cases infected with pre-Omicron VOCs, reflecting the relatively low occurrence of breakthrough infections before the Omicron burst.

In conclusion, vaccine-induced pre-Omicron immunity boosted by 3^rd^ dose or by breakthrough infection remains at least partially effective against all tested Omicron lineages, albeit with a substantial decrease compared to the ancestral B.1. The level of neutralization 4 months after the 3^rd^ vaccine dose is comparable to that induced by infection alone at the time of acute infection. In contrast, antibodies elicited by pre-Omicron infection alone less efficiently bind Spike determinants and as a consequence, less effectively neutralize Omicron sublineages. Thus immunity obviously differs qualitatively between infected, vaccinated and BTI, even if all from pre-Omicron exposures. This diversity is increasing further with the exponential increase in infections due to Omicron, adding more intricate layers to the immune landscape. The impact of immune imprinting on the viral landscape will have to be evaluated as it may contribute further evolutionary windows for the virus.

## Author Contributions

DPB designed the study and acquired funding; MK, GG, TS and VA attended patients, managed and selected samples; ESS and JYS characterized viral strains. ESS, JYS and DPB performed neutralization experiments. ESS and DPB curated and analyzed the data; ESS, DPB and CD drafted the manuscript. Authors had full access to all the data in the study, and they accept responsibility to submit for publication.

The funding source had no role in the writing of the manuscript or the decision to submit it for publication, nor in data collection, analysis, or interpretation nor any aspect pertinent to the study.

## Acknowledgments

This research was funded by the Fonds National de la Recherche du Luxembourg (FNR COVID-19 FT-1 (**14718697** NEUTRACOV), by the Rotary Clubs Luxembourg, and by Ministère de I’Education et de la Recherche du Luxembourg. The APC was funded by FNR.

## Data sharing

No clinical data was collected for this study. Deidentified leftover plasma samples and swabs were provided by Centre Hospitalier de Luxembourg and by Laboratoire National de Santé (LNS). The researchers involved in the study had no access to any data or identity.

## Institutional Review Board Statement

The study was subject to an internal review and was approved by the LIH Institutional Review Board (14718697 NeutraCoV) and conducted in accordance with the Declaration of Helsinki. As no regulatory issues were identified, and ethical review is not a requirement for this type of work, it was decided that a full ethical review would not be necessary. Any patient/participant/sample identifiers included were not known to anyone (e.g., research group, patients or participants themselves) outside the hospital staff who collected them, so cannot be used to identify individuals.

## Informed Consent Statement

Patient consent was waived due to the use of anonymized left-over samples for the validation of research tests in line with GPDR guidelines.

