## supplemental information for "Vaccine- and BTI-elicited pre-Omicron immunity more effectively neutralizes Omicron sublineages BA.1, BA.2, BA.4 and BA.5 than pre-Omicron infection alone"

### Supplementary Figures and legends

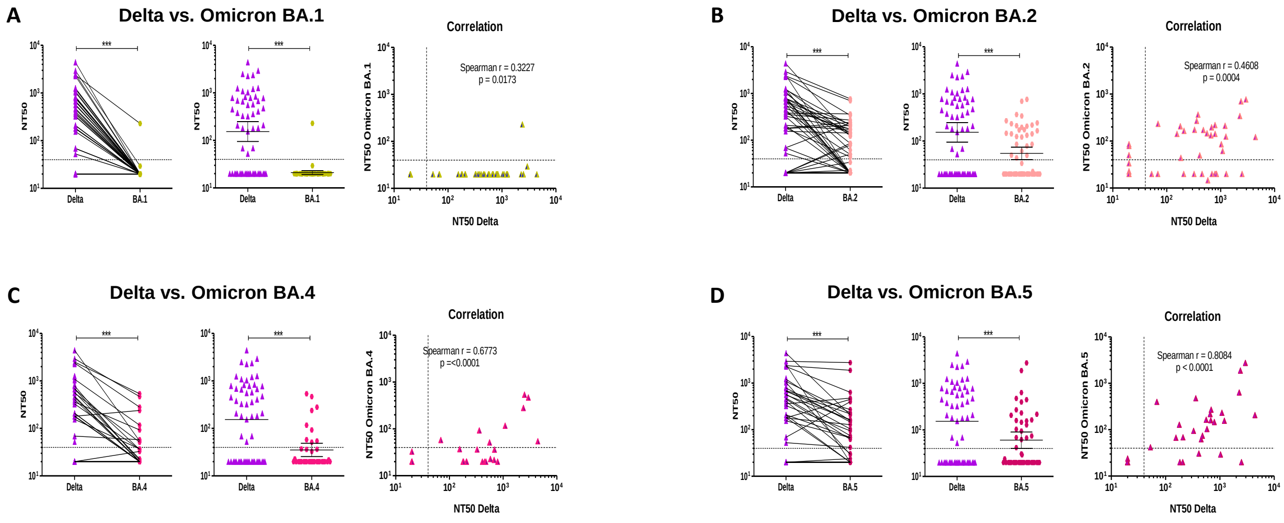


**Supplementary Figure 1. Pairwise comparison of 50% neutralizing titers (NT50) of convalescent sera against Delta and Omicron sublineages.** NT50 between Delta and Omicron sublineages BA.1 (**A**), BA.2 (**B**), BA.4 (**C**) and BA.5 were compared (**D**). Cells were infected with the strain indicated on the x-axis. Paired samples (left panel), Geometric Mean with 95% Confidence interval (middle panel) and Spearman correlation (right panel) are shown. The dotted line represents the 1:40 serum dilution cut-off. Differences between groups were compared using a Wilcoxon signed rank test. P-values < 0.05 were considered significant. *: p<0.05; **: p<0.01; ***: p<0.001.

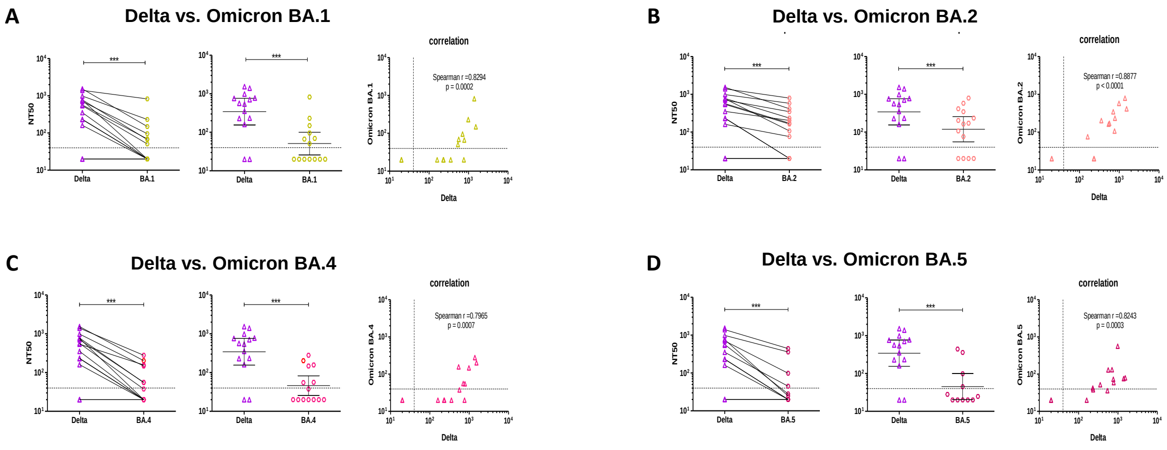


**Supplementary Figure 2. Pairwise comparison of 50% neutralizing titers (NT50) of sera from triple-vaccinated individuals against Delta and Omicron sublineages.** NT50 between Delta and Omicron sublineages BA.1 (**A**), BA.2 (**B**), BA.4 (**C**) and BA.5 were compared (**D**). Cells were infected with the strain indicated on the x-axis. Paired samples (left panel), Geometric Mean with 95% Confidence interval (middle panel) and Spearman correlation (right panel) are shown. The dotted line represents the 1:40 serum dilution cut-off. Differences between groups were compared using Wilcoxon signed rank test. P-values < 0.05 were considered significant. *: p<0.05; **: p<0.01; ***: p<0.001.

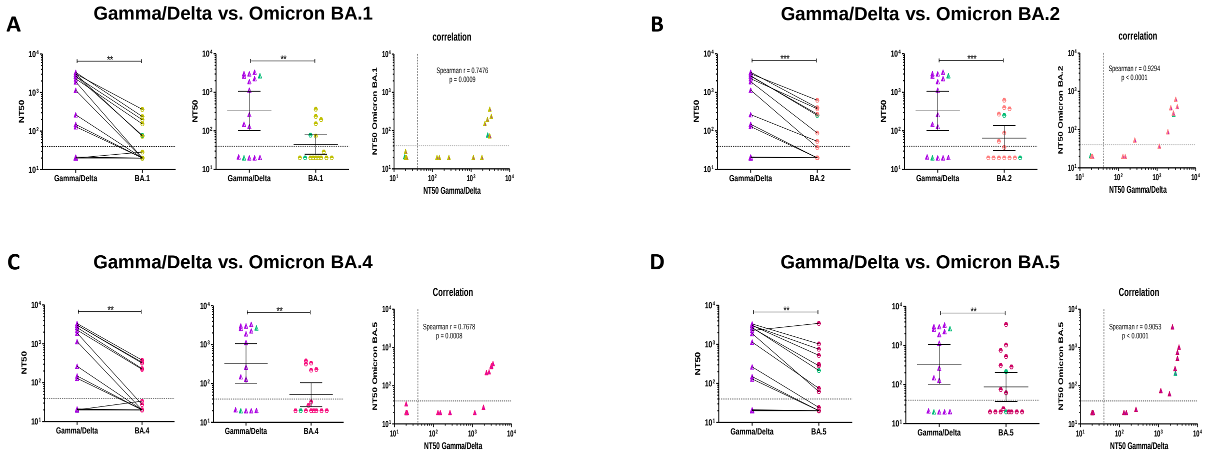


**Supplementary Figure 3. Pairwise comparison of 50% neutralizing titers (NT50) of BTI sera against Delta and Omicron sublineages.** NT50 between Delta and Omicron sublineages BA.1 (**A**), BA.2 (**B**), BA.4 (**C**) and BA.5 were compared (**D**). Cells were infected with the strain indicated on the x-axis. Paired samples (left panel), Geometric Mean with 95% Confidence interval (middle panel) and Spearman correlation (right panel) are shown. The dotted line represents the 1:40 serum dilution cut-off. Gamma-BTI are identified with green symbols. Differences between groups were compared using Wilcoxon signed rank test. P-values < 0.05 were considered significant. *: p<0.05; **: p<0.01; ***: p<0.001**.**

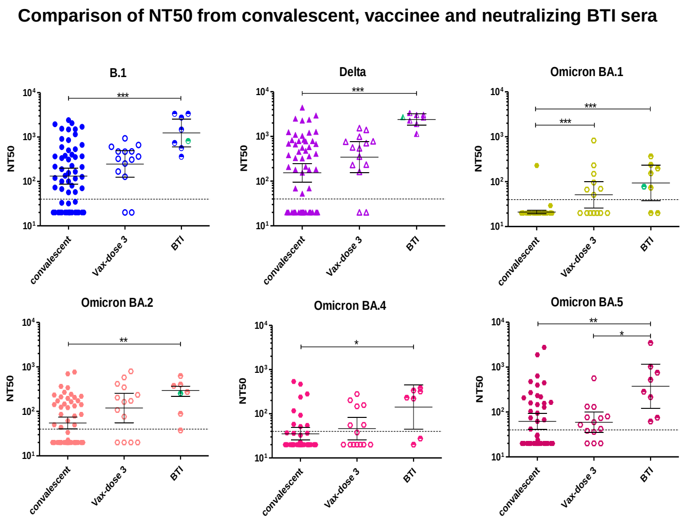


**Supplementary Figure 4. Comparison of neutralizing activities of convalescent, vaccinee and neutralizing BTI sera against B.1, Delta and Omicron BA.1, BA.2, BA.4 and BA.5.** The infecting strain is indicated above each panel. For BTI sera, the Gamma-

BTI patients infected with Gamma are represented with green symbols. Only neutralizing BTI sera are shown to highlight differences. Differences between groups were compared using a Kruskal-Wallis test followed by a Dunn's multiple comparison post-hoc test. P-values < 0.05 were considered significant. *: $p<0.05$; **: $p<0.01$; ***: $p<0.001$.
